# Whole-brain multivariate hemodynamic deconvolution for multi-echo fMRI with stability selection

**DOI:** 10.1101/2022.09.30.510299

**Authors:** Eneko Uruñuela, Javier Gonzalez-Castillo, Charles Zheng, Peter Bandettini, César Caballero-Gaudes

## Abstract

Conventionally, analysis of functional MRI (fMRI) data relies on available information about the experimental paradigm to establish hypothesized models of brain activity. However, this information can be inaccurate, incomplete or unavailable in multiple scenarios such as resting-state, naturalistic paradigms or clinical conditions. In these cases, blind estimates of neuronal-related activity can be obtained with paradigm-free analysis methods such as hemodynamic deconvolution. Yet, current formulations of the hemodynamic deconvolution problem have three important limitations: 1) their efficacy strongly depends on the appropriate selection of regularization parameters, 2) being univariate, they do not take advantage of the information present across the brain, and 3) they do not provide any measure of statistical certainty associated with each detected event. Here we propose a novel approach that addresses all these limitations. Specifically, we introduce MvME-SPFM (multivariate multi-echo sparse paradigm free mapping), a novel hemodynamic deconvolution algorithm that operates at the whole brain level and adds spatial information via a mixed-norm regularization term over all voxels. Additionally, MvME-SPFM employs a stability selection procedure that removes the need to select regularization parameters and also lets us obtain an estimate of the true probability of having a neuronal-related BOLD event at each voxel and time-point based on the area under the curve (AUC) of the stability paths. Besides, the formulation is tailored for multi-echo fMRI acquisitions, which allows us to better isolate fluctuations of BOLD origin on the basis of their linear dependence with Echo Time (TE) and to assign physiologically interpretable units (i.e., changes in the apparent transverse relaxation 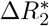) to the resulting deconvolved events. We demonstrate that this algorithm outperforms existing state-of-the-art deconvolution approaches, and shows higher spatial and temporal agreement with the activation maps and BOLD signals obtained with a standard model-based linear regression approach, even at the level of individual neuronal events. Consequently, the proposed algorithm provides more reliable estimates of neuronal-related activity, here in terms of 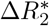, for the study of the dynamics of brain activity when no information about the timings of the BOLD events is available. This algorithm will be made publicly available as part of the *splora* Python package.

## 1. Introduction

Functional magnetic resonance imaging (fMRI) data analysis relies on the blood oxygenation level-dependent (BOLD) contrast as a proxy to localize neuronal activity and to study the functional organization of the human brain in vivo when performing a task or at rest. Due to the nature of the paradigms, task and resting state fMRI data are analyzed with different techniques. Often, the analysis of task fMRI data is performed using general linear models (GLM) that calculate statistical parametric maps of brain activity by building hypothetical timecourses of the BOLD responses to the experimental paradigm, thus exploiting the knowledge of the timings of the stimuli. However, the analysis of other types of fMRI paradigms, such as resting-state, naturalistic paradigms, or clinically-relevant assessments, cannot be performed with a GLM given that the timings of the stimuli are unknown, inaccurate or insufficient, and hence requires a paradigm free approach. Such data are typically analyzed with correlation-based methods; for example, static and dynamic functional connectivity (Preti et al., 2017), edge-centric measures (Faskowitz et al., 2020a), and inter-subject correlations (Hasson et al., 2004) Another method often used to analyze resting-state fMRI data are co-activation patterns (CAPs) (Liu et al., 2013, 2018).

However, as all these techniques operate on the BOLD signal, they are affected by the blurring that the hemodynamic response introduces to the signal, which makes the interpretation of the analyses uncertain. In order to undo this blurring effect and obtain more reliable estimates of the neuronal activity, various deconvolution techniques can be used (Glover, 1999; Gitelman et al., 2003; Gaudes et al., 2010, 2012, 2013; Caballero-Gaudes et al., 2019; Hernandez-Garcia & Ulfarsson, 2011; Karahanoglu et al., 2013; Cherkaoui et al., 2019; Costantini et al., 2022; Hütel et al., 2021). These techniques are able to blindly (i.e., with no information about the timings of neuronal events) estimate the neuronal activity that induces the BOLD response by assuming a hemodynamic response function (HRF) and solving an inverse problem with additional constraints to overcome the ill-posed nature of hemodynamic deconvolution (Uruñuela et al., 2021a).

The ability of deconvolution algorithms to estimate neuronal activity in a paradigm-free manner has been exploited in a number of applications. For instance, deconvolution techniques have been used on resting-state fMRI data to explore time-varying activity (Petridou et al., 2013; Karahanoğlu & Ville, 2015; Preti et al., 2017; Keilholz et al., 2017; Lurie et al., 2020; Bolton et al., 2020), to decode the flow of spontaneous thoughts and actions across different cognitive and sensory domains (Tan et al., 2017; Gonzalez-Castillo et al., 2019), and to investigate modulatory interactions within and between resting-state functional networks (Di & Biswal, 2013). These methods have also been applied in clinical conditions to detect the foci of interictal events in epilepsy patients without the use of EEG recordings (Lopes et al., 2012; Karahanoglu et al., 2013; Tobias et al., 2022), to investigate functional dissociations found during non-rapid eye movement sleep associated with reduced consolidation of information and impaired consciousness (Tarun et al., 2021), and to detect functional signatures of prodromal psychotic symptoms and anxiety at rest in patients with schizophrenia (Zöller et al., 2019).

Despite the range of deconvolution methods that have been developed, few capitalize on the various properties of fMRI data, such as the advantages of multi-echo fMRI for denoising fMRI data (Bright & Murphy, 2013; Kundu et al., 2017), or the use of tissue-based or parcellation-based information to improve the accuracy of the estimates of neuronal activity. Recent exceptions include deconvolution algorithms that incorporate a multivariate formulation to perform spatio-temporal deconvolution (Bolton et al., 2019a; Uruñuela et al., 2021b; Costantini et al., 2022). In addition, one deconvolution algorithm has been presented that exploits the mono-exponential decay model of the multi-echo fMRI signal: multi-echo sparse paradigm free mapping (ME-SPFM) (Caballero-Gaudes et al., 2019). Furthermore, approaches have been developed to estimate the likelihood of having a neuronal event at each time-point and for each voxel by means of logistic regression (Bush & Cisler, 2013; Bush et al., 2015) or Gaussian mixture models (Pidnebesna et al., 2019). Wouldn’ t it be nice to obtain a measure of the probability of each voxel containing a neuronal event at each time-point for regularized estimators while exploiting the mono-expontial decay and spatio-temporal properties of the multi-echo fMRI signal?

In this work, we propose a novel approach for the hemodynamic deconvolution of multi-echo fMRI data that operates at the whole-brain level (i.e., multivariate formulation) to incorporate spatial information through a mixed-norm regularization term. Furthermore, we propose a stability selection procedure (Meinshausen & Bühlmann, 2010) that makes the estimation of the neuronal activity more robust to the selection of the regularization parameters, while providing the likelihood of having a neuronal-related event at each time-point and for each voxel. Using multi-echo fMRI data acquired from 10 healthy subjects (16 datasets) we demonstrate that the proposed multivariate multi-echo paradigm free mapping (MvME-SPFM) algorithm not only provides more robust estimates of the neuronal activity, but also yields a measure of the probability of each voxel containing a neuronal event at each time-point. Moreover, MvME-SPFM returns quantitative estimates of 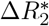 in interpretable units (s^-1^), which is relevant for functional analysis across different acquisition methods and field strengths.

## 2. Theory

### 2.1. Voxel-wise signal model for multi-echo paradigm free mapping

The analysis of BOLD fMRI data usually assumes that the signal *y*(*t*) acquired for a voxel *v* is described by the convolution between the activity-inducing signal *s*(*t*) driving the BOLD response and the hemodynamic response *h*(*t*) itself (Boynton et al., 1996; Glover, 1999), plus an additional term *e*(*t*) representing noise. Considering that the signal measured by the scanner is sampled at every TR seconds, the acquired signal can be written in discrete form as: 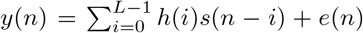, for *n* = 1,…, *N*, where *N* is the number of observations in the time-series, and *L* is the discrete-time length of the hemodynamic response function (HRF).

Hence, the signal model can be written in matrix notation as:

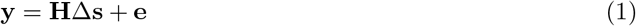

where **y**, 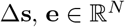 are the voxel’s time-series, the activity-inducing signal changes and the noise term, respectively, and 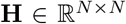 is the Toeplitz convolution matrix defined by the HRF (Gitelman et al., 2003; Gaudes et al., 2013).

For gradient-echo fMRI acquisitions, the voxel’s time-series in terms of the signal percentage change has a linear relationship with the echo time (TE) as 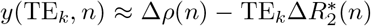, where 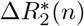 denotes the BOLD-like signal changes and Δ*ρ*(*n*) corresponds to changes in the net magnetization, for instance due to head motion (Kundu et al., 2017). The signal changes associated to fluctuations in the net magnetization can be effectively reduced in preprocessing, for example using multi-echo independent component analysis (Kundu et al., 2012; Caballero-Gaudes et al., 2019), anbd are neglected hereinafter. Hence, considering that neuronal-related signal changes Δ**s** produce a change in 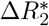, the signal model in Eq.(1) can be adapted to contain the signal acquired at all *K* echo-times (TE) via concatenation:

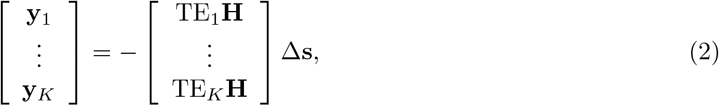

which can be simplified into 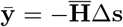. An estimate of the activity that induces the BOLD response ŝ can be obtained by solving an ordinary least-squares problem such as:

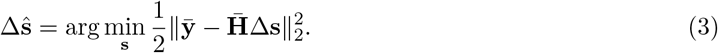

However, solving the equation above is an ill-posed problem given the high collinearity of the convolution matrix **H** due to the overlap between shifted HRFs, which introduces large variability in the estimates of **s**.

In practice, this excess of variability can be reduced by introducing additional assumptions about the activity-inducing signal in the form of regularization terms. For instance, we could assume that the activity-inducing signal is well represented by a reduced subset of non-zero coefficients at the fMRI timescale that trigger the BOLD responses. This assumption can be mathematically represented with a sparsity-promoting regularization term such as the *ℓ*_1_-norm that is added to the data fitting term in Eq.(3) (Tibshirani, 1996; Gaudes et al., 2013).

Hence, the activity-inducing signal in a single voxel can be blindly detected from the multi-echo signals by solving the following inverse problem (Caballero-Gaudes et al., 2019):

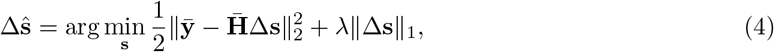

where *λ* is the regularization parameter that regulates the level of sparsity of the estimates given the *ℓ*_1_-norm, which is defined as 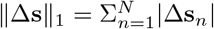. The tuning of the regularization parameter is challenging and requires the careful selection of an adequate value in order to avoid overfitting (i.e., false detection of the activity-inducing signal) or underfitting (i.e., no detection of the activity-inducing signal).

### 2.2. Whole-brain signal model for multi-echo paradigm free mapping

Assuming that the shape of the hemodynamic response can be similarly modeled across all brain voxels, the previous voxel-wise (i.e., univariate) model in Eq.(2) can be extended straightforwardly to a multivariate formulation that considers all the voxels *V* of the brain:

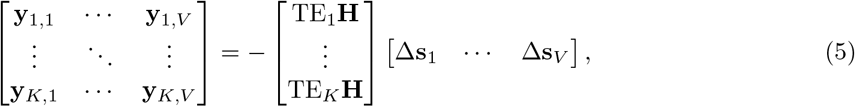

which can be simplified into 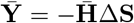, where 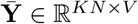, 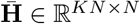 and 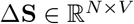.

The inverse problem in Eq.(4) can be directly adapted to be solved at the whole-brain using the multivariate formulation in Eq.(5). More interestingly though, solving the inverse problem at the wholebrain level opens up many possibilities in the form of additional regularization terms to take advantage of the spatial information for an informed estimation of the activity-inducing signal Δ**Ŝ**. For instance, mixed-norms in the form of *ℓ_p,q_* can be employed to separate coefficients into groups that are blind to each other, while the coefficients within a group are treated together (Kowalski, 2009). Hence, regularization terms based on mixed-norms can promote spatio-temporal structures that are observed in fMRI signals.

Here, we add an *ℓ*_2,1_ + *ℓ*_1_ mixed-norm regularization term (Gramfort et al., 2011) to the multivariate convex problem to promote the co-activation of the activity-inducing signal ΔŜ considering the coefficients of the voxels of the brain (*V*) at time *n* as one group:

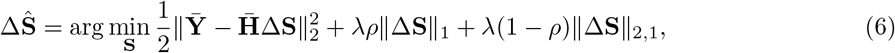

where *ℓ*_2,1_-norm is defined as 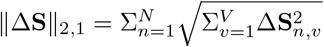, and 0 < *ρ* < 1 is a parameter that controls the tradeoff between the sparsity introduced by the *ℓ*_1_-norm and the grouping of voxels promoted by the *ℓ*_2,1_-norm so that the estimation of one voxel coefficient at time *n* is influenced by the estimates of the rest of the brain voxels at the same time. Note that when *ρ* = 1 Eq. (6) is the whole-brain equivalent of Eq. (4) On the other hand, the regularization parameter *λ* can be adapted voxel-wise in order to account for differences in the signal-to-noise ratio across voxels. Consequently, the multivariate deconvolution problem can be written as:

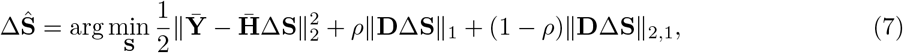

where 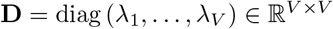 is a diagonal matrix with the voxel-specific values of *λ*. In practice, a criterion must be used to select the voxel-specific *λ*s. Instead, we propose the use of stability selection to avoid this critical choice (see Section 3.2).

Therefore, given the convex nature of the inverse problem in Eq. (7), estimates of Δ**Ŝ** can be calculated using the fast iterative shrinkage-thresholding algorithm (FISTA) (Beck & Teboulle, 2009) with the following proximity operator for *ℓ*_1_ + *ℓ*_2,1_:

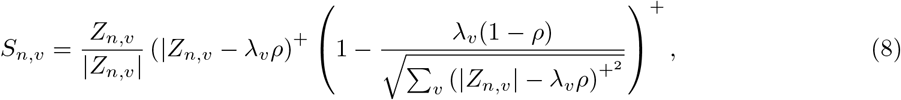

where 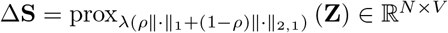, (*x*)^+^ = max (*x*, 0) for 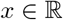, and 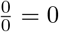 by convention.

## 3. Methods

### 3.1. fMRI data acquisition and preprocessing

The evaluation of the proposed MvME-SPFM was performed on ME-fMRI data acquired in 10 subjects using a multi-task rapid event-related paradigm. Six subjects performed two functional runs, the other 4 subjects only performed 1 run due to scanning time constraints (i.e., a total of 16 datasets). All participants gave informed consent in compliance with the NIH Combined Neuroscience International Review Board-approved protocol 93-M-1070 in Bethesda, MD. A thorough description of the MRI acquisition protocols and experimental tasks in the experimental design can be found in (Gonzalez-Castillo et al., 2016), only those details that are relevant to this analysis are given here.

MRI data was acquired on a General Electric 3T 750 MRI scanner with a 32-channel receive-only head coil (General Electric, Waukesha, WI). Functional scans were acquired with a ME gradient-recalled echoplanar imaging (GRE-EPI) sequence (flip angle = 70° for 9 subjects, flip angle = 60° for 1 subject, TEs = 16.3/32.2/48.1 ms, TR = 2 s, 30 axial slices, slice thickness = 4 mm, in-plane resolution = 3 × 3 mm^2^, FOV 192 mm, acceleration factor 2, number of acquisitions = 220). Functional data was acquired with ascending sequential slice acquisitions, except in one subject where the acquisitions were interleaved. In addition, high resolution T1-weighted MPRAGE and proton density images were acquired per subject for anatomical alignment and visualization purposes (176 axial slices, voxel size = 1 × 1 × 1 mm^3^, image matrix = 256 × 256).

Each run of data acquisition consisted of 6 trials with 5 different tasks each: biological motion observation (BMOT), finger tapping (FTAP), passive viewing of houses (HOUS), listening to music (MUSI), and sentence reading (READ). We refer the reader to that paper for details on the preprocessing steps, and comparison with alternative single-echo models for deconvolution. This data had previously been employed, preprocessed and ME-ICA denoised for the evaluation of the ME-SPFM algorithm in (Caballero-Gaudes et al., 2019).

### 3.2. Stability selection and the regularization parameter *λ*

The choice of the regularization parameter λ is crucial to obtain accurate estimates of Δ**Ŝ**. Although the value of *λ* of each voxel could be fixed ad-hoc, previous work has opted for the use of model selection criteria, such as the Bayesian Information Criterion (BIC), on the regularization path (Caballero-Gaudes et al., 2019), computed by means of the least angle regression (LARS) algorithm (Efron et al., 2004). Even though the use of BIC performed well for ME-SPFM (Caballero-Gaudes et al., 2019) and its single-echo counterpart (SPFM) (Gaudes et al., 2013), due to its high specificity, it can be problematic for certain voxels where the BIC curve might present multiple local minima or even fail to present a clear minima for the evaluated range of *λ*.

In this work, we propose a more robust procedure to address this shortcoming with the usage of the stability selection method (Meinshausen & Bühlmann, 2010). Moreover, the stability selection procedure presented here yields the probability to have a non-zero coefficient in the activity-inducing signal at each time-point. Specifically, our implementation of the stability selection procedure generates *T* = 30 surrogates by randomly subsampling 60% of the time-points (we also tested a more computationally expensive version with *T* = 100 surrogates that yielded very similar results). The convolution matrix **H** is subsampled accordingly. The subsampled data is then employed to solve the inverse problem in Eq. (6) for a range of different values of *λ* using the fast iterative shrinkage thresholding algorithm (FISTA). For each voxel, we select a logarithmically spaced sequence of 30 values between 5% and 95% of the voxel-specific maximum *λ* possible to more accurately sample the lower range. Then, for each time-point and value of *λ*, stability selection calculates the ratio (probability) of surrogates where the estimated coefficient at each time-point is non-zero. As illustrated in Figure 1, these probabilities build the so-called stability paths, which resemble the well-known regularization paths of conventional regularized estimators (e.g., LASSO, Ridge Regression) that plot the amplitude of the coefficients for each *λ*.

**Figure 1:**
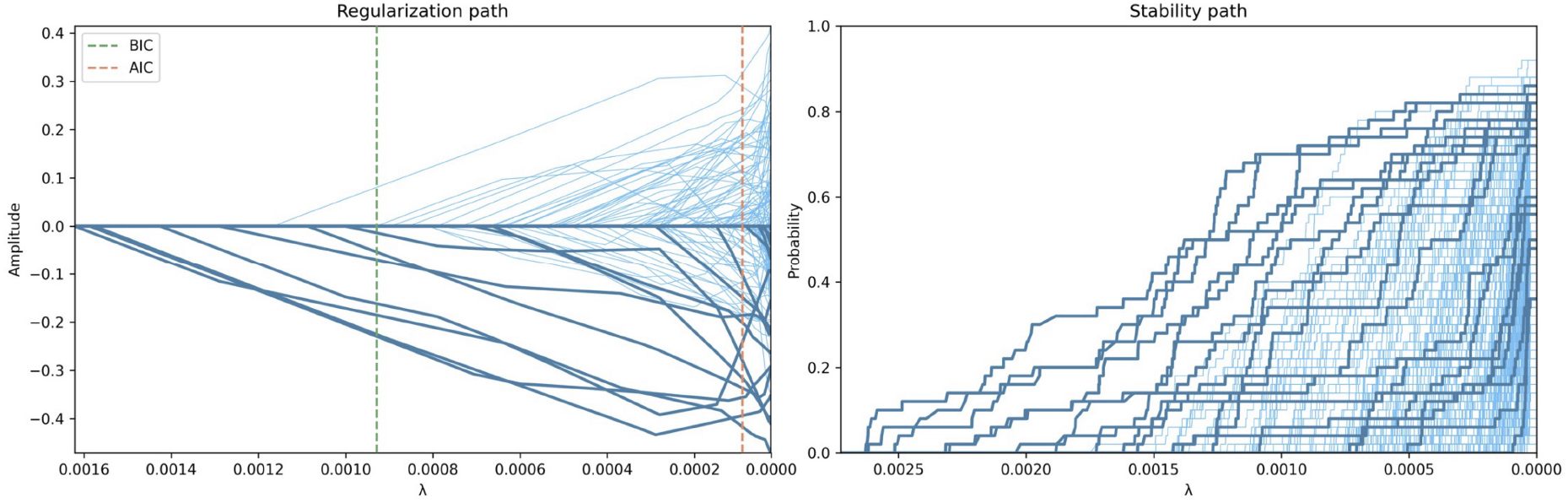
Example of the regularization path and the stability path for a voxel timeseries with *ρ* = 1. On the left, the regularization path shows the amplitude of each coefficient estimate Δ**Ŝ** (one per TR). At first, all the coefficients are zero and successively they become non-zero as *λ* decreases towards zero, which corresponds to the orthogonal least squares solution (i.e., no regularization). On the right, the corresponding stability path plots the probablity that each coefficient estimate is non-zero for each value of *λ* based on the stability selection procedure. Note that both paths can have a different maximum value of *λ* given the subsampling step in the stability selection. The darker lines denote the coefficient estimates corresponding to the TRs during the task-related events.

Unlike the original stability selection procedure, which sets a given probability threshold to select the final set of non-zero coefficients (Meinshausen & Bühlmann, 2010), we calculate the area under the curve (AUC) of the stability path of each time-point as an index of confidence of having a non-zero coefficient across the evaluated range of *λ*. As a result, the AUC timecourse provides a measure of the probability of having neuronal-related activity at each time-point and voxel. Next, the AUC time-series are thresholded according to the histogram of AUC values in a region of non-interest (hereinafter, denoted as the null AUC histogram) to yield a sparse representation of the signal. Alternatively, a null distribution of AUC values could be generated from surrogate data (Liégeois et al., 2021). Accordingly, when employing stability selection, the individual voxels’ estimates might not be equivalent to the voxels’ estimates in any single one of the whole-brain models that can be forumulated with a given value of *λ* in Eq.(6) or **D** in Eq. (7)), but are rather obtained by computing area-under-the-curve (AUC) values for neuronal-related events.

Finally, we apply a fitting step to each voxel by defining a reduced convolution model with the selected non-zero coefficients and fitting it by means of a conventional orthogonal least squares estimator. This step reduces the bias towards zero imposed by the sparsity-promoting regularization terms, and thus obtains more realistic estimates of the neuronal-related signal (here, in terms of 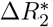) (Caballero-Gaudes et al., 2019).

### 3.3. Balancing the spatial regularization

The *ℓ*_2,1_-norm regularization term in Eq. (6) promotes structured spatio-temporal sparsity in the sense that the estimates of all brain voxels at a given time-point are treated as a group and this term forms a constraint on the number of groups with at least one non-zero estimate to model the data. Assuming that *ρ* = 0, either the value of all the voxel estimates at one time point can be non-zero or all of them are nulled. Hence, this regularization term considers spatial information from all brain voxels for the deconvolution since the value of a given voxel coefficient also depends on the rest of the voxels.

To illustrate the effect of the corresponding regularization parameter *ρ*, in this work we solve the multivariate regularization problem in Eq. (6) using stability selection for *ρ* = 1, *ρ* = 0.5 and *ρ* = 0.; i.e., applying the sparsity-promoting *ℓ*_1_-norm only, equally weighting the sparsity and spatial regularizations, and employing the *ℓ*_2,1_-norm spatial regularization only, respectively.

### 3.4. Comparison with conventional timing-based GLM analyses

To evaluate how the multivariate formulation combined with stability selection improves the accuracy of the estimates of Δ**Ŝ** compared with its univariate counterpart ME-SPFM using the BIC for voxel-wise selection of *λ* (Caballero-Gaudes et al., 2019), we calculated the spatial sensitivity, specificity and overlap (using a Dice coefficient metric) of the MvME-SPFM activation maps using the trial-level GLM-based activation maps (*p* ≤ 0.05) as the ground truth. These GLM-based maps were obtained from the optimally combined and ME-ICA denoised data, and only negative 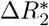 (i.e., Δ**Ŝ** < 0 that generate a positive BOLD response) were considered for the computation of the Dice coefficients.

For the MvME-SPFM, we considered the following two strategies for thresholding the AUC timeseries in order to define the corresponding activation maps:

- **Static thresholding:** The estimates of Δ**Ŝ** obtained with the novel MvME-SPFM technique that utilizes stability selection, where the AUC threshold was chosen as the 95th percentile of the histogram of AUC in deep white matter voxels (i.e., a fixed, static threshold), which were labeled after tissue segmentation of the T1-weighted anatomical MRI using *3dSeg* in AFNI, and eroding 4 voxels of the resulting white matter tissue mask at anatomical resolution.
- **Time-depending thresholding:** The estimates of Δ**Ŝ** obtained with the novel MvME-SPFM technique with stability selection, where the AUC threshold varies temporally according to the 95th percentile of the null histogram of AUC at each time-point. This implementation was based on the hypothesis that a time-dependent (TD) threshold would be able to better control for widespread spurious deconvolved changes in Δ**Ŝ**, for instance due to head motion or deep breaths.

## 4. Results

The output of deconvolution algorithms such as ME-SPFM and the proposed MvME-SPFM is a 4D dataset that matches the dimensions (both spatial and temporal) of the input data, i.e., it is a movie of the estimated 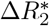 maps. In addition, the use of stability selection generates the area under the curve (AUC) 4D output dataset, which indicates the probability of having a neuronal-related event at each time-point for every voxel in the brain.

Figure 2 depicts the area under the curve (AUC) time-series and maps obtained with stability selection for *ρ* = 0.5 in representative voxels of each task in the paradigm (indicated with a cross in the maps), where the AUC maps correspond to single time-points signaled by the blue arrows. The AUC time-series of the ST and TD thresholding approaches are shown on top of the original AUC time-series. The AUC maps depict spatial patterns of 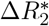 where regions that are typically involved in the tasks show higher probabilities of having neuronal-related activity compared with other brain regions.

**Figure 2:**
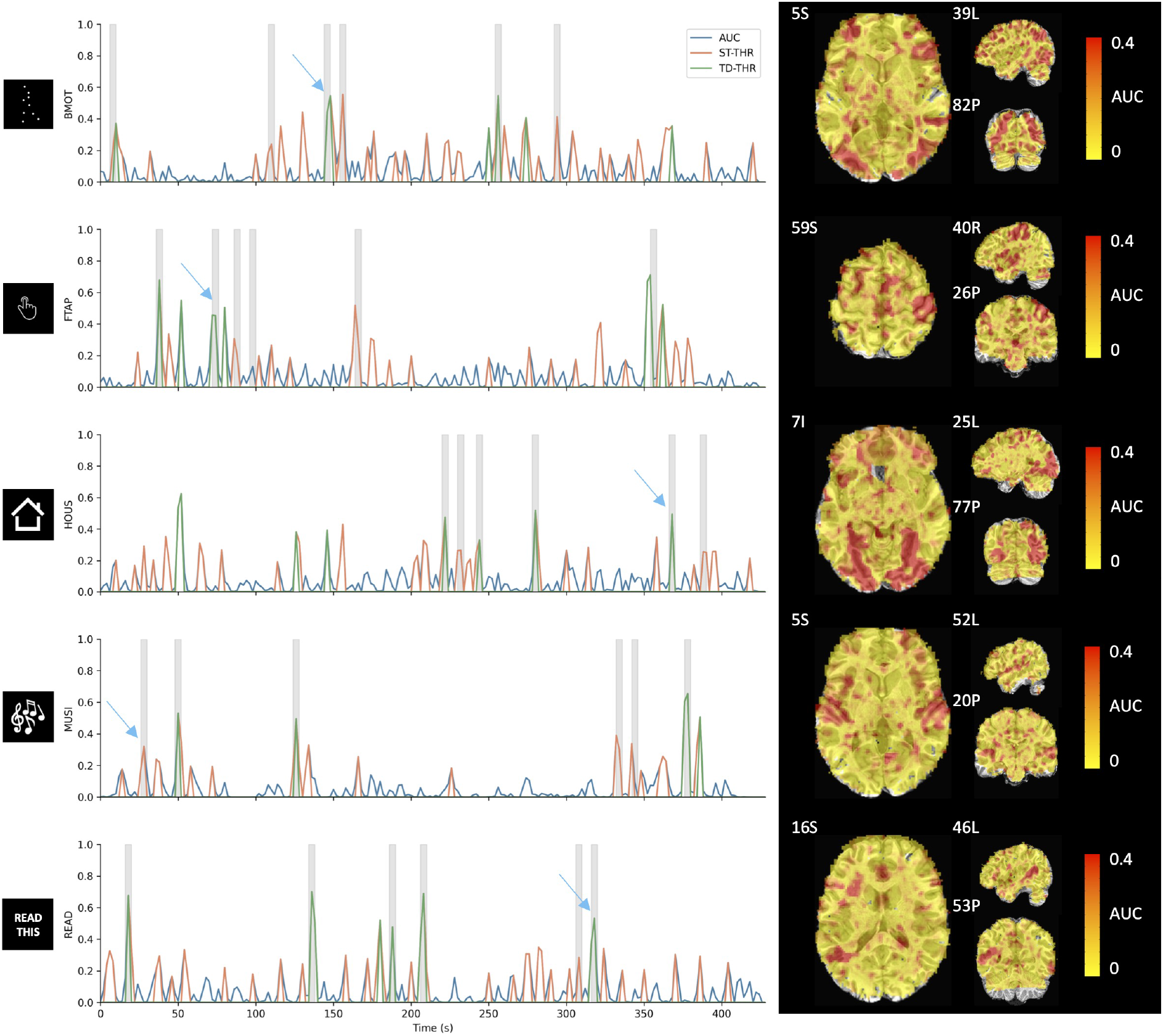
**Left:** Original (blue), ST thresholded (orange) and TD thresholded (green) AUC time-series for a representative voxel for each task in the paradigm (*ρ* = 0.5). Note that the three time-series are overlaid; i.e., the static and time-dependent time-courses are thresholded versions of the original AUC. Gray blocks depict the onset and duration of each trial. **Right:** AUC maps at the time-points signaled by the blue arrows.

Figure 3 displays the comparison of the 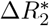 maps obtained by solving the inverse problem in Eq. (6) with a fixed selection of *λ* (1^st^ row) and with the use of stability selection (2^*nd*^, 3^*rd*^ and 4^*th*^ rows) for *ρ* = {0,0.5,1}. The 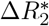 maps obtained with a fixed selection of *λ* equal to the noise estimate of the first echo volume (1^st^ row) are very sensitive to the selection of *ρ*. Similar observations were obtained with other values of *λ*. With a selection of *ρ* = 1, only the *ℓ*_1_-norm regularization term is applied, which produces 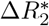 maps with few non-zero coefficients. With *ρ* = 0, only the *ℓ*_2,1_-norm spatial regularization is applied, which yields a 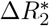 map that covers the entire brain and does not exhibit a spatial pattern in concordance with the task. However, a selection of *ρ* = 0.5 yields a 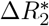 map that is more similar to the activity maps often observed when participants are asked to look at the image of a house, depicting negative 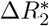 in bilateral fusiform regions. In contrast, the use of stability selection yields AUC maps (row 2) and the corresponding 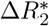 maps after each thresholding strategy (rows 3-4) reveal activation patterns concordant with those often seen for viewing houses regardless of the selection of *ρ*. In other words, the 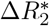 maps obtained with stability selection are less sensitive to the selection of *ρ* while obviating the need to choose *λ*. In fact, the spatial correlations between the AUC maps for each pair of *ρ*’s were nearly equal to 1 for all time points (average correlations are 0.97 between *ρ* = {0,0.5}, 0.98 between *ρ* = {0,1}, and 0.97 between *ρ* = {0.5,1}). In addition, it can be seen that using a TD threshold yields BOLD signal changes that are more confined to the expected areas in bilateral fusiform cortices than the ST threshold. Due to the high similarity of the AUC maps for any value of *ρ*, only the results for *ρ* = 0.5 are discussed hereinafter.

**Figure 3:**
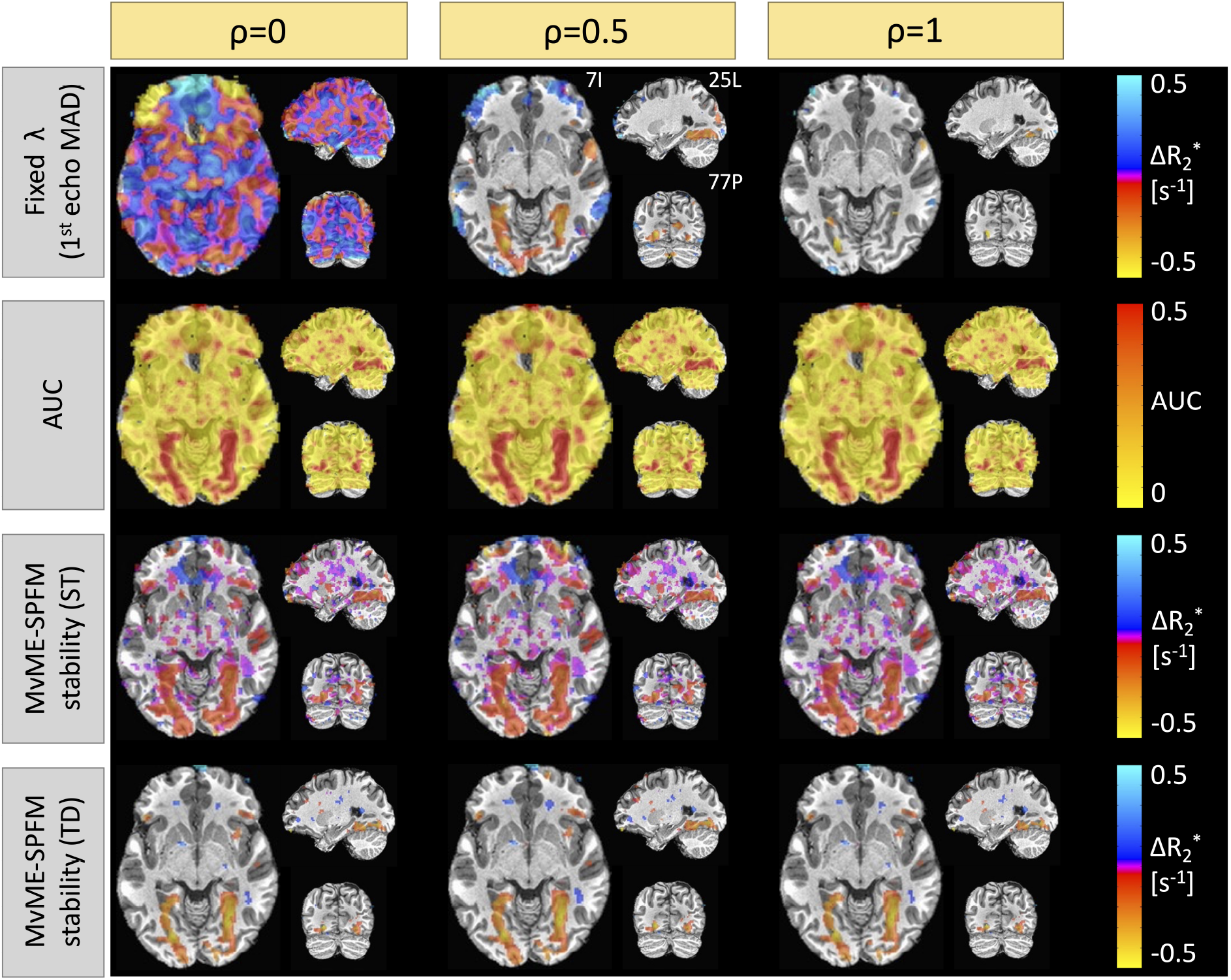
Comparison of the 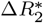 maps obtained with a fixed selection of *λ* (row 1) and the use of stability selection (rows 2-4: AUC, stability selection with static thresholding (ST), and stability selection with time-dependent thresholding (TD)) for *ρ* = 0 (column 1), *ρ* = 0.5 (column 2), and *ρ* = 1 (column 3). These maps correspond to a single-trial event of the house-viewing task (HOUS).

Figure 4 provides an in-depth view of how the time-dependent thresholding operates when motion- and respiration-related artifacts are present in the data. The grayplot (Power, 2017) in Figure 4A clearly shows bands spanning throughout the entire brain that illustrate significant changes in the amplitude of the signal. The source of these signal changes can be attributed to head motion events (see Euclidean norm in Figure 4C) and deep breaths (see arrows for respiration volume signal (Chang et al., 2009) in Figure 4D). The respiration-related events cause a drop in the global signal (see Figure 4B) seconds after the peak in the respiration volume signal. Interestingly, our results show a decrease in the equivalent ST percentile that corresponds to the 95th TD threshold (Figure 4E) at the instances of these large respiratory-related events. This decrease can also be observed in the corresponding AUC value of the TD thresholding strategy as shown in Figure 4F. The distributions of AUC values at the time-points with respiratory- and motion-related artifacts have a shorter tail than the distribution of the AUC values at the time-points where subjects performed the task. Hence, in these events the TD thresholding strategy is able to adjust the threshold so that the final estimates of 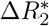 specifically capture task-activated voxels while excluding voxels that are affected by artifacts. The higher specificity of the TD thresholding strategy can be clearly seen in the ROC curves shown in Figure 4H-L. The use of stability selection with the TD threshold yields more specific estimates of 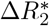 than with ST thresholding or the original ME-SPFM method, while the sensitivity is slightly reduced. On the other hand, the use of stability selection with a ST threshold improves the sensitivity of the 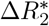 estimates compared to the original ME-SPFM technique while preserving its specificity.

**Figure 4:**
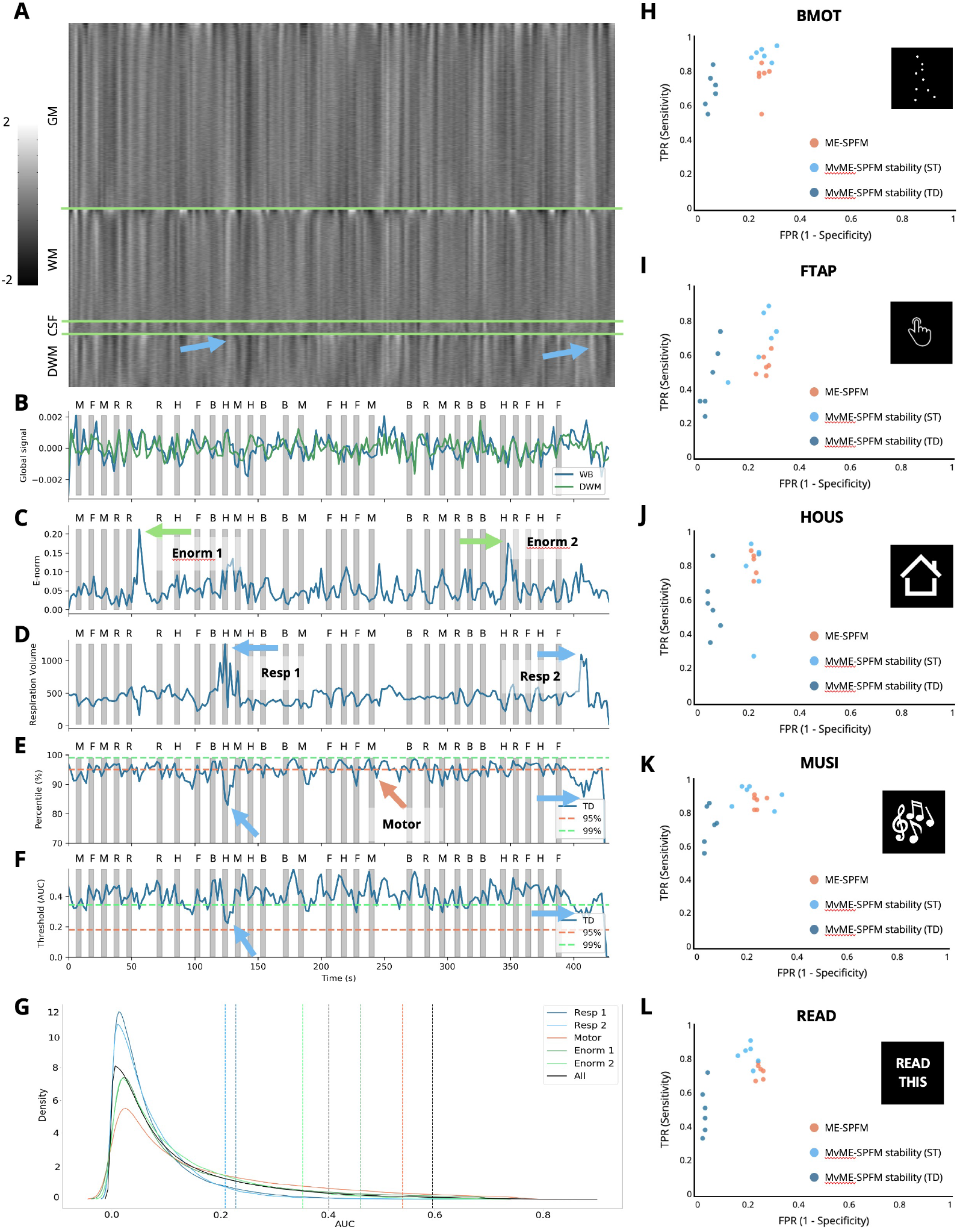
A look at the data of a representative subject with motion and respiration artifacts. **A:** Grayplot of the second echo volume. The grayplot is divided into 4 sections: gray matter (GM), white matter (WM), cerebrospinal fluid (CSF), and deep white matter (DWM). **B:** Time-series of the global signal calculated in the whole brain (WB, blue) and the deep white matter (DWM, green). **C:** Euclidean norm (e-norm) of the temporal derivative of the realignment parameters. **D:** respiration volume signal. **E:** AUC percentile corresponding to the time-dependent threshold (lines at 95th and 99th percentiles are shown for reference). **F:** AUC values corresponding to the time-dependent threshold are shown in blue. The horizontal dashed lines indicate the 95th (orange) and 99th (green) percentiles corresponding to ST thresholding. Gray bars in **B-F** indicate the onset and duration of each trial in the paradigm, with their respective initials on top. Blue arrows point out two respiration-related events, green arrows point out two motion-related events, and the orange arrow points out a finger-tapping event. **G:** Probability density functions (estimated by kernel density estimate) of the AUC values corresponding to the instances of the two respiratory-related events (blue lines), a representative time-point of one finger-tapping trial (orange line), the two largest peaks in the e-norm trace (green lines), and the overall AUC distribution (black). The corresponding coloured vertical dashed lines indicate the AUC value for the 95th percentile of the TD thresholding approach, along with the 95th and 99th AUC values of ST thresholding. **H-L:** Receiver operating characteristic (ROC) curves for the original ME-SPFM (orange), and proposed MvME-SPFM technique with the use of stability selection with the ST (light blue) and TD (dark blue) thresholding approaches for this dataset. The ROC plots depict the sensitivity and specificity of the methods at correctly estimating the activity maps that correspond to the 6 trials of each task in the paradigm.

Figure 5 illustrates the activation maps of representative single-trial events of each task for the same subject depicted in Figure 4. We compared the activation maps of the proposed MvME-SPFM formulation using the two thresholding approaches with the activation maps obtained with a single-trial GLM and the previous ME-SPFM approach. While all PFM methods exhibit activation maps that highly resemble those obtained with the single-trial GLM analysis, differences between the methods can be observed. For instance, although the use of stability selection with a ST thresholding approach yields maps with clusters of activation of comparable size and location to those found with ME-SPFM, in certain noisy trials (e.g., see HOUS-Trial 1), the ST-thresholding MvME-SPFM maps can yield reduced spatial specificity, probably related to spurious, scattered changes in 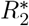. Across all tasks, the maps obtained with TD thresholding exhibit a notably larger resemblance to the single-trial GLM, showing higher spatial specificity and lower sensitivity compared to the other two PFM methods.

**Figure 5:**
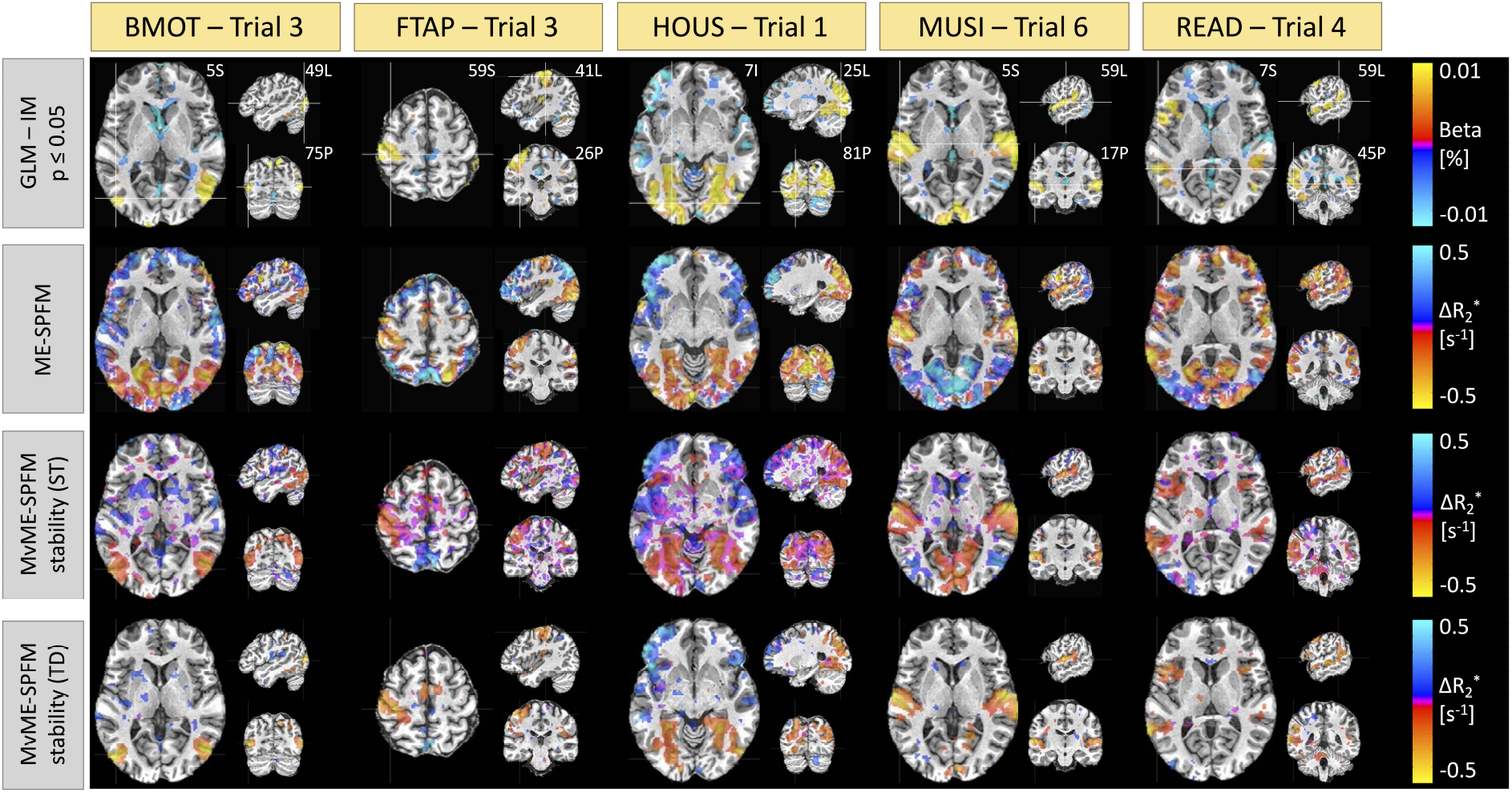
Comparison of single-trial activation maps obtained with a GLM (row 1) thresholded at *p* ≤ 0.05, the original ME-SPFM formulation with a fixed selection of *λ* (row 2), the novel MvME-SPFM technique with stability selection, *ρ* = 0.5 and a static threshold (ST, row 3), and using a time-dependent threshold (TD, row 4). A representative trial is shown for each task. All the maps correspond to the same subject and run shown in Figure 4.

Figure 6 depicts the time-series of the estimated 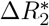 and denoised BOLD, i.e., 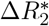 convolved with the HRF, for a representative voxel of each task for the subject depicted in Figures 4 and 5 and compared to a reference voxel in the lateral ventricles. The location of the voxels is shown in the corresponding maps in Figure 5. The ST thresholding approach detects 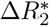 events of the activity-inducing signal that correctly match the timings of the stimuli (i.e., high temporal sensitivity), but also shows events that occur in the resting state and do not coincide with any activity-evoking trial. Based on comparison with the events detected in the time series extracted from the lateral ventricles, it can be conjectured that some of these events might be due to artifactual and physiological fluctuations that remain in the signal after preprocessing. On the other hand, 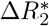 values estimated with the TD thresholding approach match the timings of the stimuli almost perfectly with few missed trials (high temporal specificity). This is supported by the few 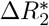 events obtained for the reference voxel in the ventricles. Likewise, the denoised BOLD time-series obtained with the TD thresholding approach clearly describes signal changes associated with the trials, whereas the denoised BOLD time-series estimated with the ST thresholding strategy fits the original data very closely, which could be interpreted as a signature of overfitting.

**Figure 6:**
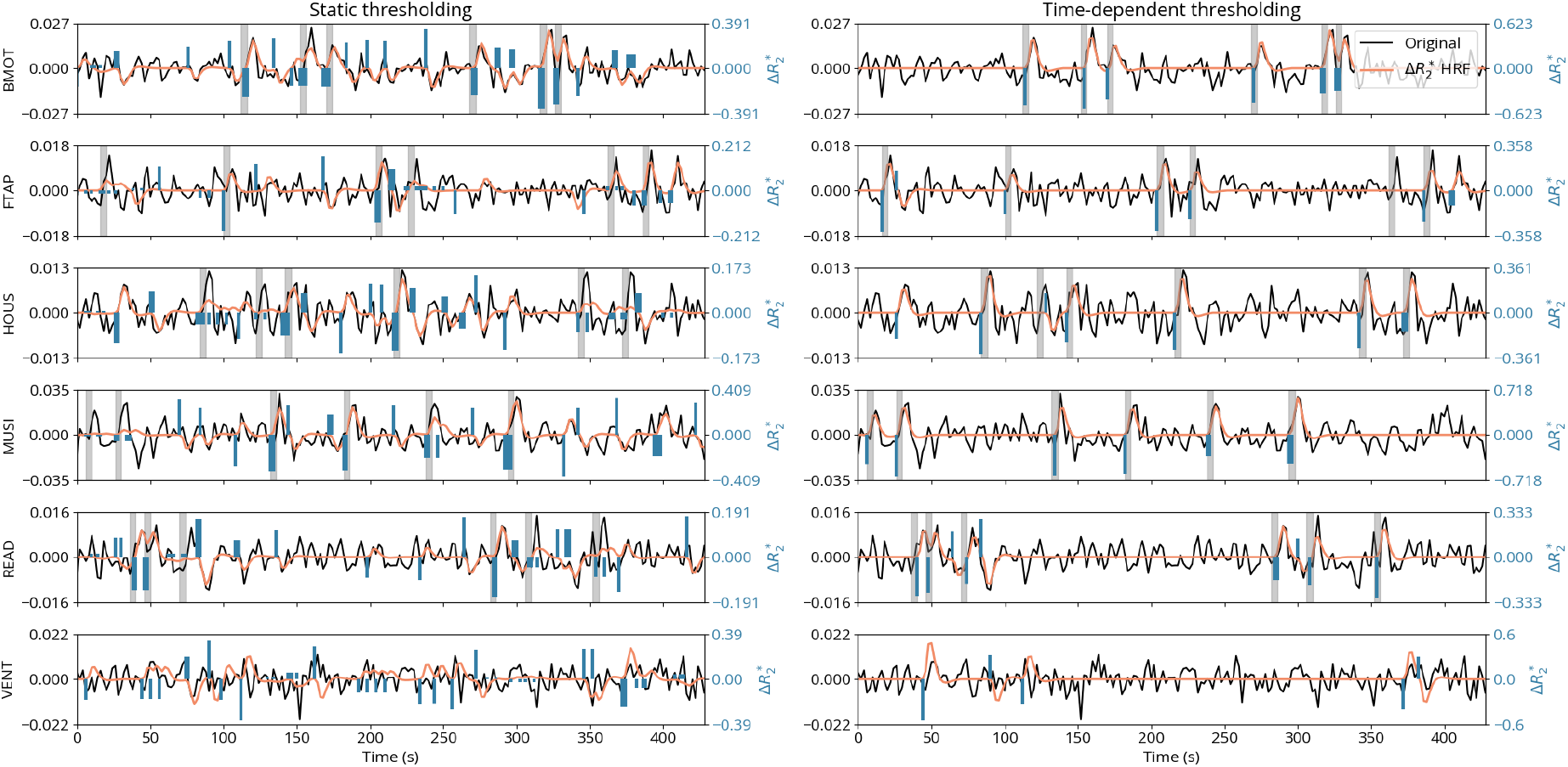
Comparison of the estimated 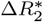 (blue) and denoised BOLD (orange), i.e., 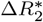 convolved with the HRF, time-series when employing the ST (left) and TD (right) thresholding approches, for representative voxels of each task (rows) as well as one voxel from the lateral ventricle for reference. The estimates shown here were obtained with *ρ* = 0.5. The preprocessed time series is shown in black. The gray bars indicate the onset and duration of each trial for each task of the experimental paradigm.

As illustrated in Figure 7, the Dice coefficient between the estimated single-trial 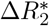 activity maps and the reference GLM activity maps (*p* ≤ 0.05) demonstrates only a slight improvement over the original ME-SPFM formulation when employing an ST thresholding approach with the novel MvME-SPFM technique. In contrast, the dice coefficients obtained with TD thresholding show a very notable increase of nearly 50% in the median of the distribution of dice coefficients compared with the original ME-SPFM approach. Similarly, the sensitivity and specificity distributions of ST thresholding demonstrate a slight improvement with respect to the original ME-SPFM formulation. On the other hand, the use of TD thresholding offers nearly perfect specificity (≥ 95%) at the cost of reduced sensitivity across all experimental conditions. Hence, increasing the specificity of the 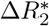 maps is more beneficial for increasing the concordance with the GLM maps than increasing the sensitivity. The receiver operating characteristic (ROC) curves in Figure 8 corroborate these observations regardless of the value of *ρ* used in the MvME-SPFM method. The estimates obtained with the ST threshold reveal an overall higher sensitivity and a slightly higher specificity compared to the original ME-SPFM technique. In contrast, the ROC curves for the TD thresholding approach show a clear improvement in specificity but lower sensitivity. These findings are in line with the results shown in Figures 3, 5 and 6, as the dice and ROC curves certify that the use of stability selection yields robust activation maps regardless of the selection of the spatial regularization term *ρ* and obviating the need to choose the temporal regularization parameter λ. An interactive version of Figures 7 and 8 is available on the GitHub repository provided in Section 7.

**Figure 7:**
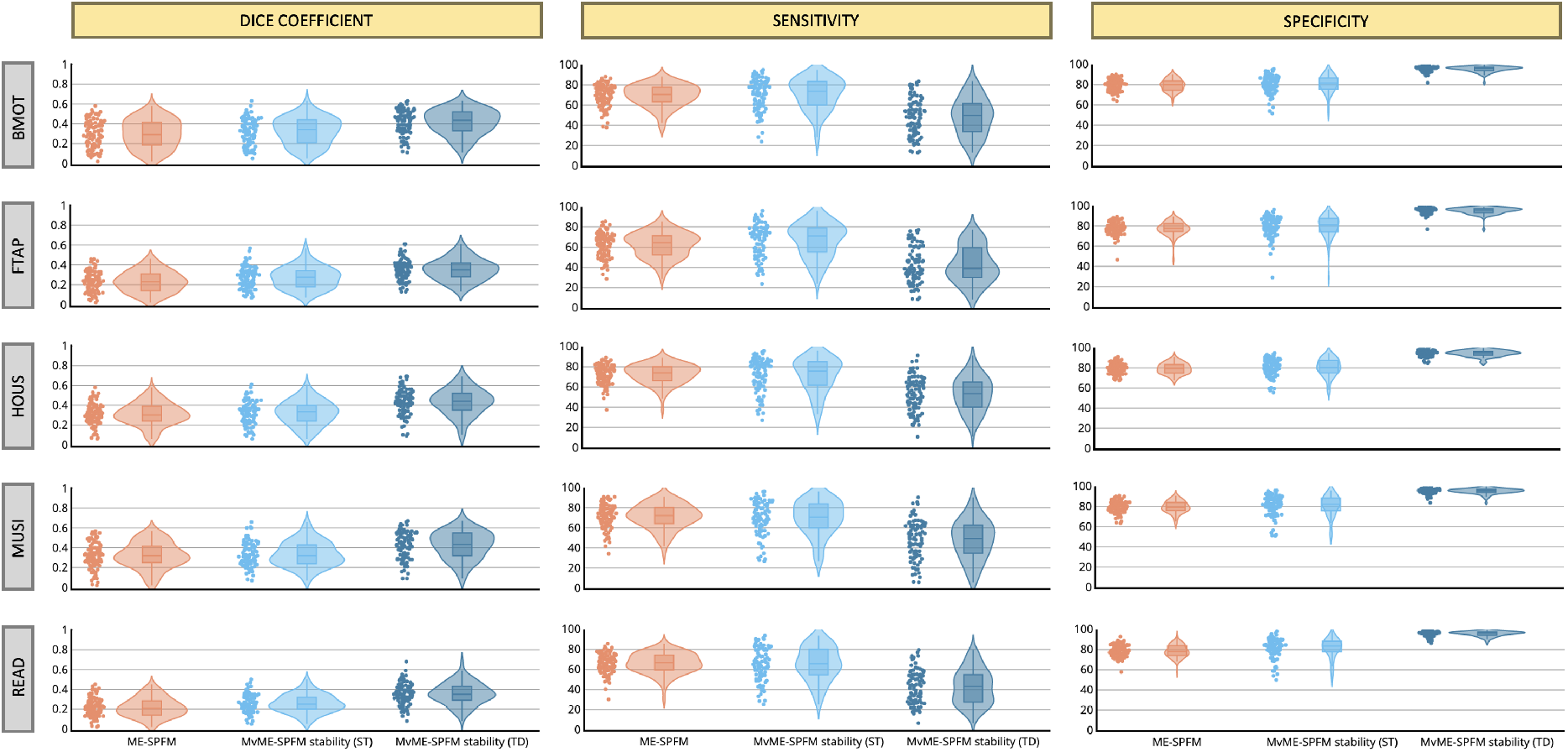
Dice coefficient (i.e., spatial overlap), sensitivity and specificity coefficients of the single-trial activation maps for each of the experimental conditions obtained with ME-SPFM, MvME-SPFM with stability selection and a static thresholding approach (ST), and MvME-SPFM with stability selection and a time-dependent thresholding approach (TD). These metrics were obtained with a selection of *ρ* = 0.5. Reference activation maps were obtained with a single trial GLM analysis and thresholded at uncorrected *p* ≤ 0.05. The density plot shows the shape of the distribution of the dice coefficients, and the box plot depicts the median with a solid line, with each box spanning from quartile 1 to quartile 3. The whiskers extend to 1.5 times the interquartile range.

**Figure 8:**
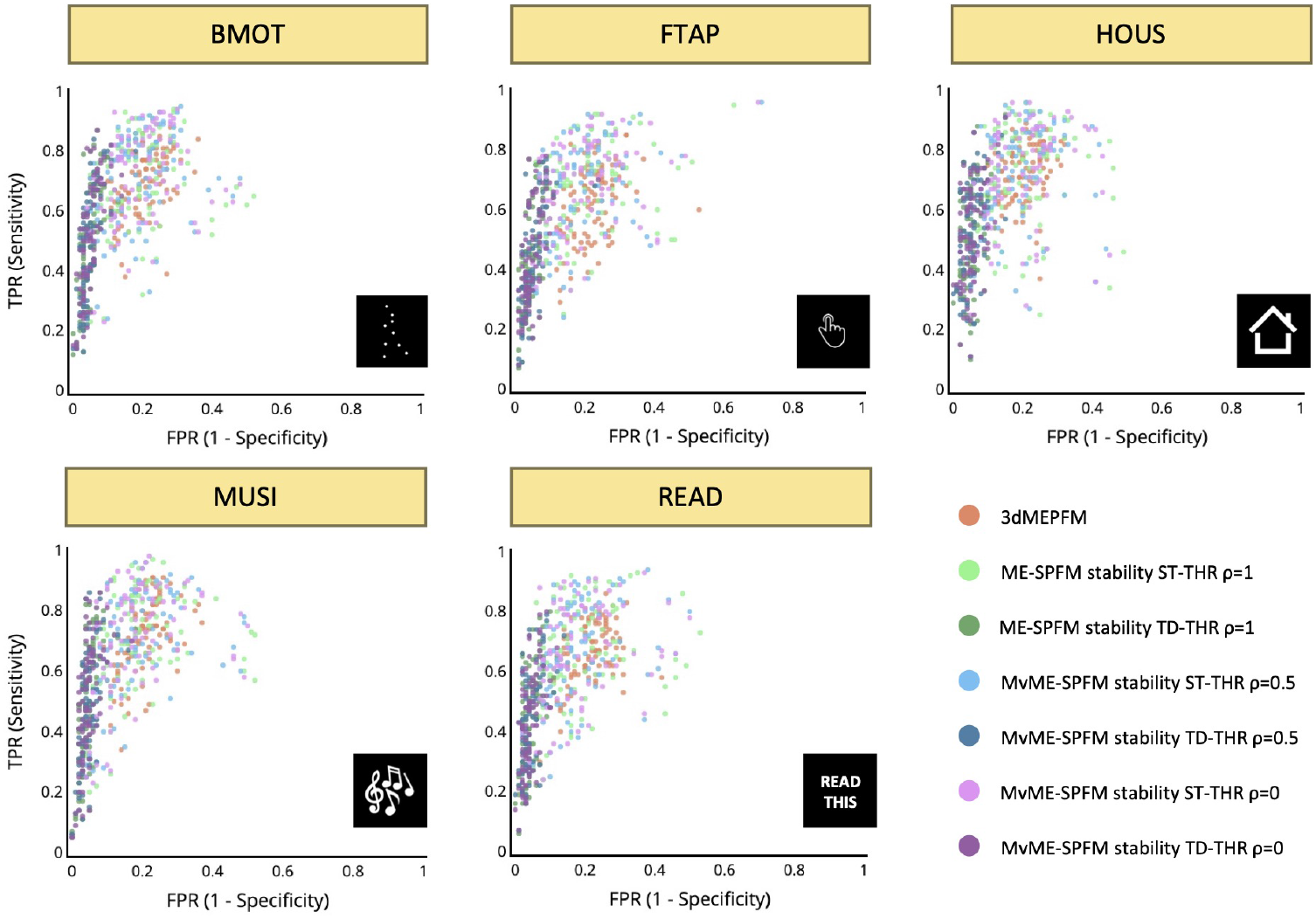
Receiver operating characteristic (ROC) curves with the sensitivity and specificity of each single trial’s activation map for all conditions and the reference map obtained with a single-trial GLM. Different colors are used for the different analyses: the original ME-SPFM, and the novel MvME-SPFM approach using stability selection with three different selections of the spatial regularization parameter *ρ* and the two different thresholding methods: static (ST) and time-dependent (TD). In each analysis each dot represents a single trial, depicting all trials across all datasets.

## 5. Discussion

The proposed whole-brain (i.e., multivariate) formulation for hemodynamic deconvolution of multiecho fMRI data with the use of stability selection achieved a closer agreement with the activation maps obtained with a single-trial GLM analysis than the original ME-SPFM method (Caballero-Gaudes et al., 2019), while obviating the need to select the temporal regularization parameter *λ* (see Figure 5). In addition, our results illustrated that the stability selection procedure also offers robustness against the choice of the spatial regularization parameter *ρ*, as the AUC maps for different selections of *ρ* were practically identical, as shown in Figure 3. Hence, although stability selection could be employed with a double selection of the regularization parameters *λ* and *ρ*, this can be avoided for computational reasons with little influence in the results. In any case, extending the proposed stability selection technique to other formulations of the hemodynamic deconvolution problem, such as the voxel-wise (i.e., univariate) single-echo (Gaudes et al., 2013; Uruñuela et al., 2020), univariate multi-echo (Caballero-Gaudes et al., 2019), or low-rank and sparse formulations (Uruñuela et al., 2021b; Cherkaoui et al., 2021), is relatively straightforward. Moreover, considering that synthesis-based models, such as Paradigm Free Mapping (Gaudes et al., 2013), and analysis-based models, such as Total Activation (Karahanoğlu et al., 2013), for temporal hemodynamic deconvolution yield identical results (Uruñuela et al., 2021a), and the fact that a multi-echo formulation provides higher accuracy for deconvolution (Caballero-Gaudes et al., 2019), we argue that the proposed MvME-SPFM method with stability selection should result in more reliable estimates of the activity-inducing signal.

One of the most interesting features of the proposed stability selection procedure is the estimation of the area under the curve (AUC) measure, which provides a new perspective for exploring fMRI data: a 4D movie with the probability of each voxel and time point containing a neuronal-related event. Therefore, the AUC time-series and maps provide complementary information to the estimates of 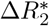, and serve as a reliability measure. Even though the AUC measures were employed here to produce the final estimates of the activity-inducing signal, they could also be exploited on their own. For instance, they could be exploited to constrain functional connectivity analysis (Tagliazucchi et al., 2016; Faskowitz et al., 2020b) to voxels and instants with a high probability of containing a neuronal-related event. Furthermore, the stability selection and the AUC metric can also be interpreted from a machine learning perspective, where the outputs from a collection of lasso learners are combined with an ensemble regression approach. Alternatively, the stability selection procedure could also be linked to Bayesian approaches where the prior is given by the range of values of the regularization term *λ* and the total posterior probability of the neuronal event is calculated as the integration of the stability paths, i.e., the AUC (see discussion in Meinshausen & Bühlmann, 2010).

Although the estimation of the AUC eliminates the need to select the spatial and temporal regularization parameters *λ* and *ρ*, it requires the use of a thresholding approach given the nature of the AUC measure, which cannot be equal to zero by definition. Here, we adopted two data-driven thresholding strategies, static (ST) and time-dependent (TD), based on the AUC values of a region where no BOLD signal changes related to neuronal activity are assumed to occur (e.g., deep white matter voxels). The use of a static AUC thresholding approach yielded higher sensitivity than the original ME-SPFM method (Caballero-Gaudes et al., 2019) while maintaining the specificity as demonstrated in Figure 8. Notably, this improvement was seen in all trials with the exception of one outlier run, regardless of the choice of the spatial regularization parameter *ρ*. Nevertheless, the use of a time-dependent thresholding approach may be even justified by the increased specificity and nearly perfect retrieval of the activity-inducing signal (row 3 in Figure 6) when motion- and respiration-related artifacts are visible in the data (see arrows in Figure 4). However, the application of the time-dependent threshold may reduce sensitivity at the single-trial level in some cases. Hence, the results shown in Figure 8 encourage the use of the static thresholding approach as an exploratory step before employing the time-dependent threshold. Other thresholding criteria could involve the comparison of AUC values obtained from surrogate (null) data (Liègeois et al., 2021) with the AUC values obtained with the original data.

Furthermore, the extension of the original ME-SPFM algorithm from a voxel-wise to a whole-brain (i.e., multivariate) regularized problem paves the way for more refined formulations that exploit the spatial characteristics and information available in fMRI and complementary imaging data into the spatial regularization term in order to improve the estimation of 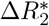. For instance, the spatial regularization could be constrained within brain regions delineated by commonly used parcellations (e.g., the Schaefer-Yeo atlas) (Karahanoğlu et al., 2013) or within neighbouring gray matter voxels (Farouj et al., 2017). Moreover, the multivariate formulation could exploit complimentary multimodal information such as structural connectvity from diffusion-based MRI data (Bolton et al., 2019b). In addition, the proposed formulation can be easily adapted to model the changes in neuronal activty in terms of its innovations, which can be more appropiate to capture sustained BOLD events (Uruñuela et al., 2021a).

Similar to the results obtained with ME-SPFM (Caballero-Gaudes et al., 2019), we observed that MvME-SPFM also detects hemodynamic events with physiologically plausible 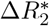 and relatively high AUC values in periods between trials when the subjects are not engaged in any activity-evoking task, whereas analysis approaches that model events with known timings (e.g., GLM) cannot find these spontaneous events. Consequently, MvME-SPFM can provide robust estimates of the activity that drives BOLD responses ocurring in spontaneous brain fluctuations (Finn et al., 2015; Tanner et al., 2022), to study individual differences in naturalistic paradigms (Finn et al., 2020), to blindly decode the subject’s engagement in a particular cognitive process from the activation maps (Poldrack, 2011; Poldrack & Yarkoni, 2016; Gonzalez-Castillo et al., 2019; Tan et al., 2017), or in clinical conditions such as the study of the urge-to-tic in patients with Tourette’s syndrome (Jackson et al., 2020).

One limitation of the proposed MvME-SPFM technique is the assumption of a particular shape of the hemodynamic response to construct the HRF matrix for deconvolution in Eq. (5). The proposed model does not account for the variability in the temporal characteristics of the HRF across the brain, which originates from differences in stimulus intensity and patterns, short inter-event intervals, or differences in the HRF shape between resting-state and task-based paradigms (Yeşilyurt et al., 2008; Sadaghiani et al., 2009; Chen et al., 2021; Polimeni & Lewis, 2021). To resolve this issue, given that the performance of MvME-SPFM is not time-locked to the trials, the current formulation could be extended to account for variability in the onset of the activity-inducing signal, as well as to introduce flexibility in the model, by employing multiple basis functions (Gaudes et al., 2012). Finally, the computational demands involved in the stability selection procedure, which solves the regularization problem in Eq. (6) for a range of *λ* values on a number of subsampled surrogate datasets, are higher than solving the regularization path and finding an adequate solution via model selection criteria as in ME-SPFM (Caballero-Gaudes et al., 2019).

## 6. Conclusion

In summary, this work proposes a new approach (MvME-SPFM) for the deconvolution of multi-echo fMRI data that exploits spatial information of the fMRI data with a whole-brain (i.e., multivariate) formulation and an *ℓ*_2,1_-norm, and yields more robust estimates of changes in 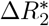 through the use of the stability selection procedure. Moreover, this work introduced a novel metric based on the area under the curve (AUC) of the stability paths that depicts the probability of having neuronal-related events at each voxel and time-point. We demonstrated that the proposed approach yields more robust and superior estimates of 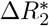 compared with the original ME-SPFM approach, and shows high spatial and temporal agreement with activation maps obtained with a GLM, while having no information about the timings of the BOLD events.

## Supporting information

Reason for updating the manuscript

## 7. Code and data availability

The code and materials used to generate the figures in this work can be found in the following GitHub repository: https://github.com/eurunuela/MvMEPFM_figures.

The Python package is available as part of *splora* in the following GitHub repository: https://github.com/eurunuela/splora.

## 8. Acknowledgements

This research was funded by the Spanish Ministry of Economy and Competitiveness (RYC-2017-21845), the Basque Government (BERC 2018-2021, PIB_2019_104, PRE_2020_2_0227), and the Spanish Ministry of Science, Innovation and Universities (PID2019-105520GB-100). This research was also possible thanks to the support of the National Institute of Mental Health Intramural Research Program (ZIAMH002783, ZICMH002968).

## 9. CRediT

Eneko Uruñuela: Conceptualization, Methodology, Software, Formal Analysis, Investigation, Writing (OD), Writing (RE), Visualization, Funding Acquisition. Javier Gonzalez-Castillo: Data Curation, Writing (RE). Charles Zheng: Writing (RE). Peter Bandettini: Funding Acquisition, Writing (RE). César Caballero-Gaudes: Conceptualization, Methodology, Software, Formal Analysis, Investigation, Data Curation, Writing (OD), Writing (RE), Visualization, Funding Acquisition.

## References

Beck, A., & Teboulle, M. (2009). A fast iterative shrinkage-thresholding algorithm for linear inverse problems. SIAM Journal on Imaging Sciences, 2, 183–202. doi:10.1137/080716542.

Bolton, T. A., Farouj, Y., Inan, M., & Ville, D. V. D. (2019a). Structurally-informed deconvolution of functional magnetic resonance imaging data. In 2019 IEEE 16th International Symposium on Biomedical Imaging (ISBI 2019). IEEE. doi:10.1109/isbi.2019.8759218.

Bolton, T. A., Farouj, Y., Inan, M., & Ville, D. V. D. (2019b). Structurally-informed deconvolution of functional magnetic resonance imaging data. In 2019 IEEE 16th International Symposium on Biomedical Imaging (ISBI 2019). IEEE. doi:10.1109/isbi.2019.8759218.

Bolton, T. A., Morgenroth, E., Preti, M. G., & Ville, D. V. D. (2020). Tapping into multi-faceted human behavior and psychopathology using fMRI brain dynamics. Trends in Neurosciences, 43, 667–680. doi:10.1016/j.tins.2020.06.005.

Boynton, G. M., Engel, S. A., Glover, G. H., & Heeger, D. J. (1996). Linear systems analysis of functional magnetic resonance imaging in human v1. The Journal of Neuroscience, 16, 4207–4221. doi:10.1523/jneurosci.16-13-04207.1996.

Bright, M. G., & Murphy, K. (2013). Removing motion and physiological artifacts from intrinsic BOLD fluctuations using short echo data. NeuroImage, 64, 526–537. doi:10.1016/j.neuroimage.2012.09.043.

Bush, K., & Cisler, J. (2013). Decoding neural events from fMRI BOLD signal: A comparison of existing approaches and development of a new algorithm. Magnetic Resonance Imaging, 31, 976–989. doi:10.1016/j.mri.2013.03.015.

Bush, K., Cisler, J., Bian, J., Hazaroglu, G., Hazaroglu, O., & Kilts, C. (2015). Improving the precision of fMRI BOLD signal deconvolution with implications for connectivity analysis. Magnetic Resonance Imaging, 33, 1314–1323. doi:10.1016/j.mri.2015.07.007.

Caballero-Gaudes, C., Moia, S., Panwar, P., Bandettini, P. A., & Gonzalez-Castillo, J. (2019). A deconvolution algorithm for multi-echo functional MRI: Multi-echo sparse paradigm free mapping. NeuroImage, 202, 116081. doi:10.1016/j.neuroimage.2019.116081.

Chang, C., Cunningham, J. P., & Glover, G. H. (2009). Influence of heart rate on the BOLD signal: The cardiac response function. NeuroImage, 44, 857–869. doi:10.1016/j.neuroimage.2008.09.029.

Chen, J. E., Glover, G. H., Fultz, N. E., Rosen, B. R., Polimeni, J. R., & Lewis, L. D. (2021). Investigating mechanisms of fast BOLD responses: The effects of stimulus intensity and of spatial heterogeneity of hemodynamics. NeuroImage, 245, 118658. doi:10.1016/j.neuroimage.2021.118658.

Cherkaoui, H., Moreau, T., Halimi, A., & Ciuciu, P. (2019). Sparsity-based blind deconvolution of neural activation signal in FMRI. In ICASSP 2019 - 2019 IEEE International Conference on Acoustics, Speech and Signal Processing (ICASSP). IEEE. doi:10.1109/icassp.2019.8683358.

Cherkaoui, H., Moreau, T., Halimi, A., Leroy, C., & Ciuciu, P. (2021). Multivariate semi-blind deconvolution of fMRI time series. NeuroImage, 241, 118418. doi:10.1016/j.neuroimage.2021.118418.

Costantini, I., Deriche, R., & Deslauriers-Gauthier, S. (2022). An anisotropic 4d filtering approach to recover brain activation from paradigm-free functional MRI data. Frontiers in Neuroimaging, 1. doi:10.3389/fnimg.2022.815423.

Di, X., & Biswal, B. B. (2013). Modulatory interactions of resting-state brain functional connectivity. PLoS ONE, 8, e71163. doi:10.1371/journal.pone.0071163.

Efron, B., Hastie, T., Johnstone, I., & Tibshirani, R. (2004). Least angle regression. The Annals of Statistics, 32. doi:10.1214/009053604000000067.

Farouj, Y., Karahanoglu, F. I., & Ville, D. V. D. (2017). Regularized spatiotemporal deconvolution of fMRI data using gray-matter constrained total variation. In 2017 IEEE 14th International Symposium on Biomedical Imaging (ISBI 2017). IEEE. doi:10.1109/isbi.2017.7950563.

Faskowitz, J., Esfahlani, F. Z., Jo, Y., Sporns, O., & Betzel, R. F. (2020a). Edge-centric functional network representations of human cerebral cortex reveal overlapping system-level architecture. Nature Neuroscience, 23, 1644–1654. doi:10.1038/s41593-020-00719-y.

Faskowitz, J., Esfahlani, F. Z., Jo, Y., Sporns, O., & Betzel, R. F. (2020b). Edge-centric functional network representations of human cerebral cortex reveal overlapping system-level architecture. Nature Neuroscience, 23, 1644–1654. doi:10.1038/s41593-020-00719-y.

Finn, E. S., Glerean, E., Khojandi, A. Y., Nielson, D., Molfese, P. J., Handwerker, D. A., & Bandettini, P. A. (2020). Idiosynchrony: From shared responses to individual differences during naturalistic neuroimaging. NeuroImage, 215, 116828. doi:10.1016/j.neuroimage.2020.116828.

Finn, E. S., Shen, X., Scheinost, D., Rosenberg, M. D., Huang, J., Chun, M. M., Papademetris, X., & Constable, R. T. (2015). Functional connectome fingerprinting: identifying individuals using patterns of brain connectivity. Nature Neuroscience, 18, 1664–1671. doi:10.1038/nn.4135.

Gaudes, C. C., Karahanoglu, F. I., Lazeyras, F., & Ville, D. V. D. (2012). Structured sparse deconvolution for paradigm free mapping of functional MRI data. In 2012 9th IEEE International Symposium on Biomedical Imaging (ISBI). IEEE. doi:10.1109/isbi.2012.6235549.

Gaudes, C. C., Petridou, N., Dryden, I. L., Bai, L., Francis, S. T., & Gowland, P. A. (2010). Detection and characterization of single-trial fMRI bold responses: Paradigm free mapping. Human Brain Mapping, 32, 1400–1418. doi:10.1002/hbm.21116.

Gaudes, C. C., Petridou, N., Francis, S. T., Dryden, I. L., & Gowland, P. A. (2013). Paradigm free mapping with sparse regression automatically detects single-trial functional magnetic resonance imaging blood oxygenation level dependent responses. Human Brain Mapping, (pp. n/a–n/a). doi:10.1002/hbm.21452.

Gitelman, D. R., Penny, W. D., Ashburner, J., & Friston, K. J. (2003). Modeling regional and psychophysiologic interactions in fMRI: the importance of hemodynamic deconvolution. NeuroImage, 19, 200–207. doi:10.1016/s1053-8119(03)00058-2.

Glover, G. H. (1999). Deconvolution of impulse response in event-related BOLD fMRI1. NeuroImage, 9, 416–429. doi:10.1006/nimg.1998.0419.

Gonzalez-Castillo, J., Caballero-Gaudes, C., Topolski, N., Handwerker, D. A., Pereira, F., & Bandettini, P. A. (2019). Imaging the spontaneous flow of thought: Distinct periods of cognition contribute to dynamic functional connectivity during rest. NeuroImage, 202, 116129. doi:10.1016/j.neuroimage.2019.116129.

Gonzalez-Castillo, J., Panwar, P., Buchanan, L. C., Caballero-Gaudes, C., Handwerker, D. A., Jangraw, D. C., Zachariou, V., Inati, S., Roopchansingh, V., Derbyshire, J. A., & Bandettini, P. A. (2016). Evaluation of multi-echo ICA denoising for task based fMRI studies: Block designs, rapid event-related designs, and cardiac-gated fMRI. NeuroImage, 141, 452–468. doi:10.1016/j.neuroimage.2016.07.049.

Gramfort, A., Strohmeier, D., Haueisen, J., Hamalainen, M., & Kowalski, M. (2011). Functional brain imaging with m/EEG using structured sparsity in time-frequency dictionaries. In Lecture Notes in Computer Science (pp. 600–611). Springer Berlin Heidelberg. doi:10.1007/978-3-642-22092-0_49.

Hasson, U., Nir, Y., Levy, I., Fuhrmann, G., & Malach, R. (2004). Intersubject synchronization of cortical activity during natural vision. Science, 303, 1634–1640. doi:10.1126/science.1089506.

Hernandez-Garcia, L., & Ulfarsson, M. O. (2011). Neuronal event detection in fMRI time series using iterative deconvolution techniques. Magnetic Resonance Imaging, 29, 353–364. doi:10.1016/j.mri.2010.10.012.

Hütel, M., Antonelli, M., Melbourne, A., & Ourselin, S. (2021). Hemodynamic matrix factorization for functional magnetic resonance imaging. NeuroImage, 231, 117814. doi:10.1016/j.neuroimage.2021.117814.

Jackson, S. R., Loayza, J., Crighton, M., Sigurdsson, H. P., Dyke, K., & Jackson, G. M. (2020). The role of the insula in the generation of motor tics and the experience of the premonitory urge-to-tic in tourette syndrome. Cortex, 126, 119–133. doi:10.1016/j.cortex.2019.12.021.

Karahanoğlu, F. I., Caballero-Gaudes, C., Lazeyras, F., & Ville, D. V. D. (2013). Total activation: fMRI deconvolution through spatio-temporal regularization. NeuroImage, 73, 121–134. doi:10.1016/j.neuroimage.2013.01.067.

Karahanoglu, F. I., Grouiller, F., Gaudes, C. C., Seeck, M., Vulliemoz, S., & Ville, D. V. D. (2013). Spatial mapping of interictal epileptic discharges in fMRI with total activation. In 2013 IEEE 10th International Symposium on Biomedical Imaging. IEEE. doi:10.1109/isbi.2013.6556819.

Karahanoğlu, F. I., & Ville, D. V. D. (2015). Transient brain activity disentangles fMRI resting-state dynamics in terms of spatially and temporally overlapping networks. Nature Communications, 6. doi:10.1038/ncomms8751.

Keilholz, S., Caballero-Gaudes, C., Bandettini, P., Deco, G., & Calhoun, V. (2017). Time-resolved resting-state functional magnetic resonance imaging analysis: Current status, challenges, and new directions. Brain Connectivity, 7, 465–481. doi:10.1089/brain.2017.0543.

Kowalski, M. (2009). Sparse regression using mixed norms. Applied and Computational Harmonic Analysis, 27, 303–324. doi:10.1016/j.acha.2009.05.006.

Kundu, P., Inati, S. J., Evans, J. W., Luh, W.-M., & Bandettini, P. A. (2012). Differentiating BOLD and non-BOLD signals in fMRI time series using multi-echo EPI. NeuroImage, 60, 1759–1770. doi:10.1016/j.neuroimage.2011.12.028.

Kundu, P., Voon, V., Balchandani, P., Lombardo, M. V., Poser, B. A., & Bandettini, P. A. (2017). Multiecho fMRI: A review of applications in fMRI denoising and analysis of BOLD signals. NeuroImage, 154, 59–80. doi:10.1016/j.neuroimage.2017.03.033.

Liégeois, R., Yeo, B. T., & Ville, D. V. D. (2021). Interpreting null models of resting-state functional MRI dynamics: not throwing the model out with the hypothesis. NeuroImage, 243, 118518. doi:10.1016/j.neuroimage.2021.118518.

Liu, X., Chang, C., & Duyn, J. H. (2013). Decomposition of spontaneous brain activity into distinct fMRI co-activation patterns. Frontiers in Systems Neuroscience, 7. doi:10.3389/fnsys.2013.00101.

Liu, X., Zhang, N., Chang, C., & Duyn, J. H. (2018). Co-activation patterns in resting-state fMRI signals. NeuroImage, 180, 485–494. doi:10.1016/j.neuroimage.2018.01.041.

Lopes, R., Lina, J., Fahoum, F., & Gotman, J. (2012). Detection of epileptic activity in fMRI without recording the EEG. NeuroImage, 60, 1867–1879. doi:10.1016/j.neuroimage.2011.12.083.

Lurie, D. J., Kessler, D., Bassett, D. S., Betzel, R. F., Breakspear, M., Kheilholz, S., Kucyi, A., Liégeois, R., Lindquist, M. A., McIntosh, A. R., Poldrack, R. A., Shine, J. M., Thompson, W. H., Bielczyk, N. Z., Douw, L., Kraft, D., Miller, R. L., Muthuraman, M., Pasquini, L., Razi, A., Vidaurre, D., Xie, H., & Calhoun, V. D. (2020). Questions and controversies in the study of time-varying functional connectivity in resting fMRI. Network Neuroscience, 4, 30–69. doi:10.1162/netn_a_00116.

Meinshausen, N., & Bühlmann, P. (2010). Stability selection. Journal of the Royal Statistical S’ociety: Series B (Statistical Methodology), 72, 417–473. doi:10.1111/j.1467-9868.2010.00740.x.

Petridou, N., Gaudes, C. C., Dryden, I. L., Francis, S. T., & Gowland, P. A. (2013). Periods of rest in fMRI contain individual spontaneous events which are related to slowly fluctuating spontaneous activity. Human Brain Mapping, 34, 1319–1329. doi:10.1002/hbm.21513.

Pidnebesna, A., Fajnerová, I., Horáček, J., & Hlinka, J. (2019). Estimating sparse neuronal signal from hemodynamic response: the mixture components inference approach,. doi:10.1101/2019.12.19.876508.

Poldrack, R. A. (2011). Inferring mental states from neuroimaging data: From reverse inference to large-scale decoding. Neuron, 72, 692–697. doi:10.1016/j.neuron.2011.11.001.

Poldrack, R. A., & Yarkoni, T. (2016). From brain maps to cognitive ontologies: Informatics and the search for mental structure. Annual Review of Psychology, 67, 587–612. doi:10.1146/annurev-psych-122414-033729.

Polimeni, J. R., & Lewis, L. D. (2021). Imaging faster neural dynamics with fast fMRI: A need for updated models of the hemodynamic response. Progress in Neurobiology, 207, 102174. doi:10.1016/j.pneurobio.2021.102174.

Power, J. D. (2017). A simple but useful way to assess fMRI scan qualities. NeuroImage, 154, 150–158. doi:10.1016/j.neuroimage.2016.08.009.

Preti, M. G., Bolton, T. A., & Ville, D. V. D. (2017). The dynamic functional connectome: State-of-the-art and perspectives. NeuroImage, 160, 41–54. doi:10.1016/j.neuroimage.2016.12.061.

Sadaghiani, S., Uğurbil, K., & Uludağ, K. (2009). Neural activity-induced modulation of BOLD post-stimulus undershoot independent of the positive signal. Magnetic Resonance Imaging, 27, 1030–1038. doi:10.1016/j.mri.2009.04.003.

Tagliazucchi, E., Siniatchkin, M., Laufs, H., & Chialvo, D. R. (2016). The voxel-wise functional connectome can be efficiently derived from co-activations in a sparse spatio-temporal point-process. Frontiers in Neuroscience, 10. doi:10.3389/fnins.2016.00381.

Tan, F. M., Caballero-Gaudes, C., Mullinger, K. J., Cho, S.-Y., Zhang, Y., Dryden, I. L., Francis, S. T., & Gowland, P. A. (2017). Decoding fMRI events in sensorimotor motor network using sparse paradigm free mapping and activation likelihood estimates. Human Brain Mapping, 38, 5778–5794. doi:10.1002/hbm.23767.

Tanner, J. C., Faskowitz, J., Byrge, L., Kennedy, D. P., Sporns, O., & Betzel, R. F. (2022). Synchronous high-amplitude co-fluctuations of functional brain networks during movie-watching,. doi:10.1101/2022.06.30.497603.

Tarun, A., Wainstein-Andriano, D., Sterpenich, V., Bayer, L., Perogamvros, L., Solms, M., Axmacher, N., Schwartz, S., & Ville, D. V. D. (2021). NREM sleep stages specifically alter dynamical integration of large-scale brain networks. iScience, 24, 101923. doi:10.1016/j.isci.2020.101923.

Tibshirani, R. (1996). Regression shrinkage and selection via the lasso. Journal of the Royal Statistical Society: Series B (Methodological), 58, 267–288. doi:10.1111/j.2517-6161.1996.tb02080.x.

Tobias, C., Uruñuela, E., Ferrer-Gallardo, V., Goldberg, H., Engelman, C., Lowe, M., Jones, S., & Caballero-Gaudes, C. (2022). Automatic detection of spatio-temporal patterns of interictal epileptic activity with fmri. [Conference Oral Scientific Session] Joint Annual Meeting ISMRM-ESMRMB & ISMRT 31st Annual Meeting,.

Uruñuela, E., Bolton, T. A. W., Van De Ville, D., & Caballero-Gaudes, C. (2021a). Hemodynamic deconvolution demystified: Sparsity-driven regularization at work. arXiv,. doi:10.48550/ARXIV.2107.12026.

Uruñuela, E., Jones, S., Crawford, A., Shin, W., Oh, S., Lowe, M., & Caballero-Gaudes, C. (2020). Stability-based sparse paradigm free mapping algorithm for deconvolution of functional MRI data. In 2020 42nd Annual International Conference of the IEEE Engineering in Medicine & Biology Society (EMBC). IEEE. doi:10.1109/embc44109.2020.9176137.

Uruñuela, E., Moia, S., & Caballero-Gaudes, C. (2021b). A low rank and sparse paradigm free mapping algorithm for deconvolution of FMRI data. In 2021 IEEE 18th International Symposium on Biomedical Imaging (ISBI). IEEE. doi:10.1109/isbi48211.2021.9433821.

Yeşilyurt, B., Uğurbil, K., & Uludağ, K. (2008). Dynamics and nonlinearities of the BOLD response at very short stimulus durations. Magnetic Resonance Imaging, 26, 853–862. doi:10.1016/j.mri.2008.01.008.

Zöller, D., Sandini, C., Karahanoğlu, F. I., Padula, M. C., Schaer, M., Eliez, S., & Ville, D. V. D. (2019). Large-scale brain network dynamics provide a measure of psychosis and anxiety in 22q11.2 deletion syndrome. Biological Psychiatry: Cognitive Neuroscience and Neuroimaging, 4, 881–892. doi:10.1016/j.bpsc.2019.04.004.

